# A metabolic pathway for bile acid dehydroxylation by the gut microbiome

**DOI:** 10.1101/758557

**Authors:** Masanori Funabashi, Tyler L. Grove, Victoria Pascal, Yug Varma, Molly E. McFadden, Laura C. Brown, Chunjun Guo, Marnix H. Medema, Steven C. Almo, Michael A. Fischbach

## Abstract

The gut microbiota synthesize hundreds of molecules, many of which are known to impact host physiology. Among the most abundant metabolites are the secondary bile acids deoxycholic acid (DCA) and lithocholic acid (LCA), which accumulate at ~500 μM and are known to block *C. difficile* growth^1^, promote hepatocellular carcinoma^2^, and modulate host metabolism via the GPCR TGR5^3^. More broadly, DCA, LCA and their derivatives are a major component of the recirculating bile acid pool^4^; the size and composition of this pool are a target of therapies for primary biliary cholangitis and nonalcoholic steatohepatitis. Despite the clear impact of DCA and LCA on host physiology, incomplete knowledge of their biosynthetic genes and a lack of genetic tools in their native producer limit our ability to modulate secondary bile acid levels in the host. Here, we complete the pathway to DCA/LCA by assigning and characterizing enzymes for each of the steps in its reductive arm, revealing a strategy in which the A-B rings of the steroid core are transiently converted into an electron acceptor for two reductive steps carried out by Fe-S flavoenzymes. Using anaerobic in vitro reconstitution, we establish that a set of six enzymes is necessary and sufficient for the 8-step conversion of cholic acid to DCA. We then engineer the pathway into *Clostridium sporogenes*, conferring production of DCA and LCA on a non-producing commensal and demonstrating that a microbiome-derived pathway can be expressed and controlled heterologously. These data establish a complete pathway to two central components of the bile acid pool, and provide a road map for deorphaning and engineering pathways from the microbiome as a critical step toward controlling the metabolic output of the gut microbiota.

## MAIN TEXT

The human gut microbiota harbor hundreds of pathways, most of which are encoded by genes that have not yet been identified^5–8^. Their small molecule products are of interest for three reasons: (i) Most derive predominantly or exclusively from the microbiota (i.e., there is no host source), and many enter the circulation, where they can have effects on peripheral tissues and organ systems. (ii) Their concentrations are similar to or exceed that of a typical drug; for example, indoxyl sulfate can accumulate in the human host at 130 mg/day^9^. Moreover, their concentration ranges are large, typically >10-fold^10^, which could help explain microbiome-mediated biological differences among people. (iii) Of the few high-abundance molecules whose biological functions are well understood, most are ligands for a key host receptor; for example, short-chain fatty acids modulate host immune function via GPR41/GPR43^11–13^. Thus, high-abundance, microbiota-derived molecules are responsible for a remarkably broad range of phenotypes conferred on the host by bacteria.

Among these pathways, 7α-dehydroxylation of the primary bile acids cholic acid (CA) and chenodeoxycholic acid (CDCA) is particularly notable because the organisms that carry it out are present at very low abundance—an estimated 1:10^6^ in a typical gut community^14^—yet they fully process a pool of primary bile acids that is ~1 mM in concentration^15^. Therefore, the flux through this pathway must be very high in the small subset of cells in which it operates, and the low-abundance organisms in the microbiome that perform this transformation have an unusually large impact on the pool of metabolites that enters the host. This pathway’s products—deoxycholic acid (DCA) and lithocholic acid (LCA)—are the most abundant secondary bile acids in humans (up to 450-700 μM in cecal contents)^16^, and are known to be important in three biological contexts: prevention of *Clostridium difficile* outgrowth^1^, induction of hepatocellular carcinogenesis^2^, and modulation of the host metabolic and immune responses^17–19^. More broadly, DCA, LCA, and their derivatives are a major component of the recirculating bile acid pool, representing >90% of the pool in the intestine and >25% in the gallbladder^15^. These microbiome-derived bile acids are therefore central to understanding the efficacy of therapeutics that target the bile acid pool and are approved or in clinical trials for primary biliary cholangitis and nonalcoholic steatohepatitis^4^.

In pioneering work, Hylemon and coworkers showed that the gut bacterium *Clostridium scindens* VPI 12708 carries out the 7α-dehydroxylation of CA to produce DCA^20^. CA serves as an inducer of 7α-dehydroxylation, leading to the discovery of a bile-acid-induced operon (termed *bai*) containing eight genes (**Fig. 1**)^21^. The postulated pathway consists of an oxidative arm in which four electrons are removed from the 3,7-dihydroxy moiety in the A/B ring system to generate a 3-oxo-4,5-6,7-didehydro intermediate, followed by a reductive arm in which six electrons are deposited to yield the 7-dehydroxylated product^21^ (**Extended Data Fig. 1**). Through heterologous expression and characterization of individual *bai* gene products, enzymes have been attributed to each step of the oxidative arm of the pathway^22–27^, but the reductive arm of the pathway remains poorly characterized. Given that this pathway generates abundant metabolites with broad biological impact, it is notable that the set of enzymes necessary and sufficient for 7α-dehydroxylation have not yet been defined. A complete understanding of the pathway would enable efforts to control the composition of the bile acid pool by engineering the microbiome.

**Fig. 1.**
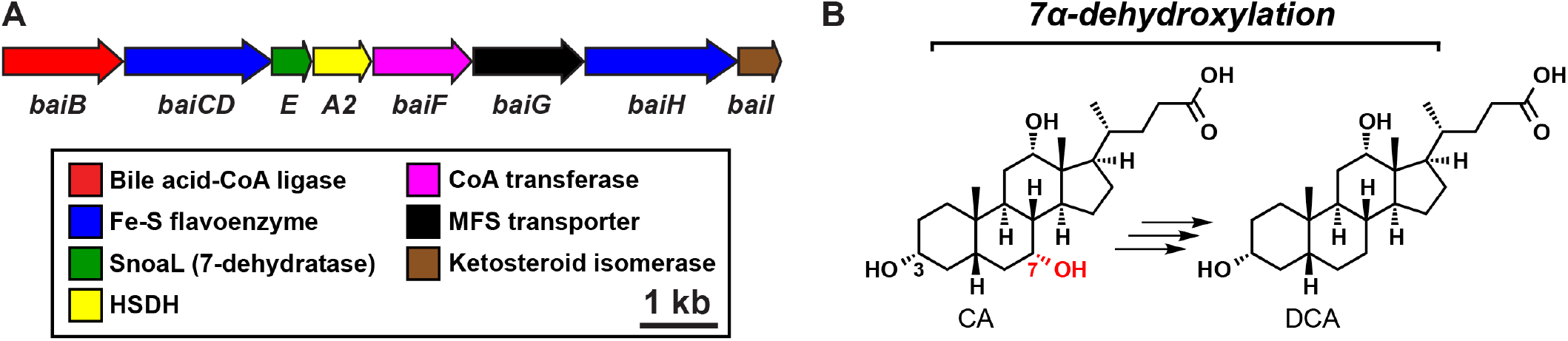
Schematic of the *bai* operon and 7α-dehydroxylation. (A) The *bai* operon consists of eight genes: seven encode enzymes and the eighth, *baiG*, encodes a transporter. It is conserved in every bacterial species known to 7α-dehydroxylate primary bile acids, and its gene products have been linked to specific steps in the pathway. (B) A simplified schematic showing the dehydroxylation of CA to DCA.

Here, by purifying and assaying pathway enzymes under anaerobic conditions, we reconstituted 7α-dehydroxylation in vitro, demonstrating that a core set of six enzymes are necessary and sufficient for the conversion of CA to DCA and revealing an unusual redox strategy in which the steroid core is transiently converted into an electron acceptor. We then transferred the pathway from its genetically intractable producer *Clostridium scindens* into *Clostridium sporogenes*, conferring production of DCA and LCA on a non-producing commensal bacterial species. These data establish a complete pathway for two central components of the bile acid pool, and they provide a genetic basis for controlling the bile acid output of the microbiome.

### Biochemical reconstitution of bile acid 7α-dehydroxylation

We first set out to de-orphan the remaining steps in the 7α-dehydroxylation pathway. Since previous studies of the *bai* enzymes involved expressing them individually in *E. coli*, we reasoned that an alternative approach in which enzymes were purified, mixed, and assayed *in vitro* could help delineate the set of enzymes necessary and sufficient for 7α-dehydroxylation. Given that the eight-gene *bai* operon is shared among all known 7α-dehydroxylating strains, we focused our efforts on the enzymes encoded by the operon. We cloned three orthologs of each enzyme, expressed them individually in *E. coli* under micro-aerobic conditions, and purified them anaerobically as N-terminal His6 fusions. Using this strategy, we obtained at least one soluble, purified ortholog of each Bai enzyme (**Extended Data Fig. 2**). When we incubated a mixture of the purified Bai enzymes with CA, NAD+, coenzyme A, and ATP under anaerobic conditions and monitored the reaction by LC-MS, we observed the time-dependent conversion of CA to DCA, indicating that the combination of BaiB, BaiCD, BaiA2, BaiE, BaiF, and BaiH is sufficient for 7α-dehydroxylation; no additional enzymes are required (**Fig. 2a, b**).

**Fig. 2.**
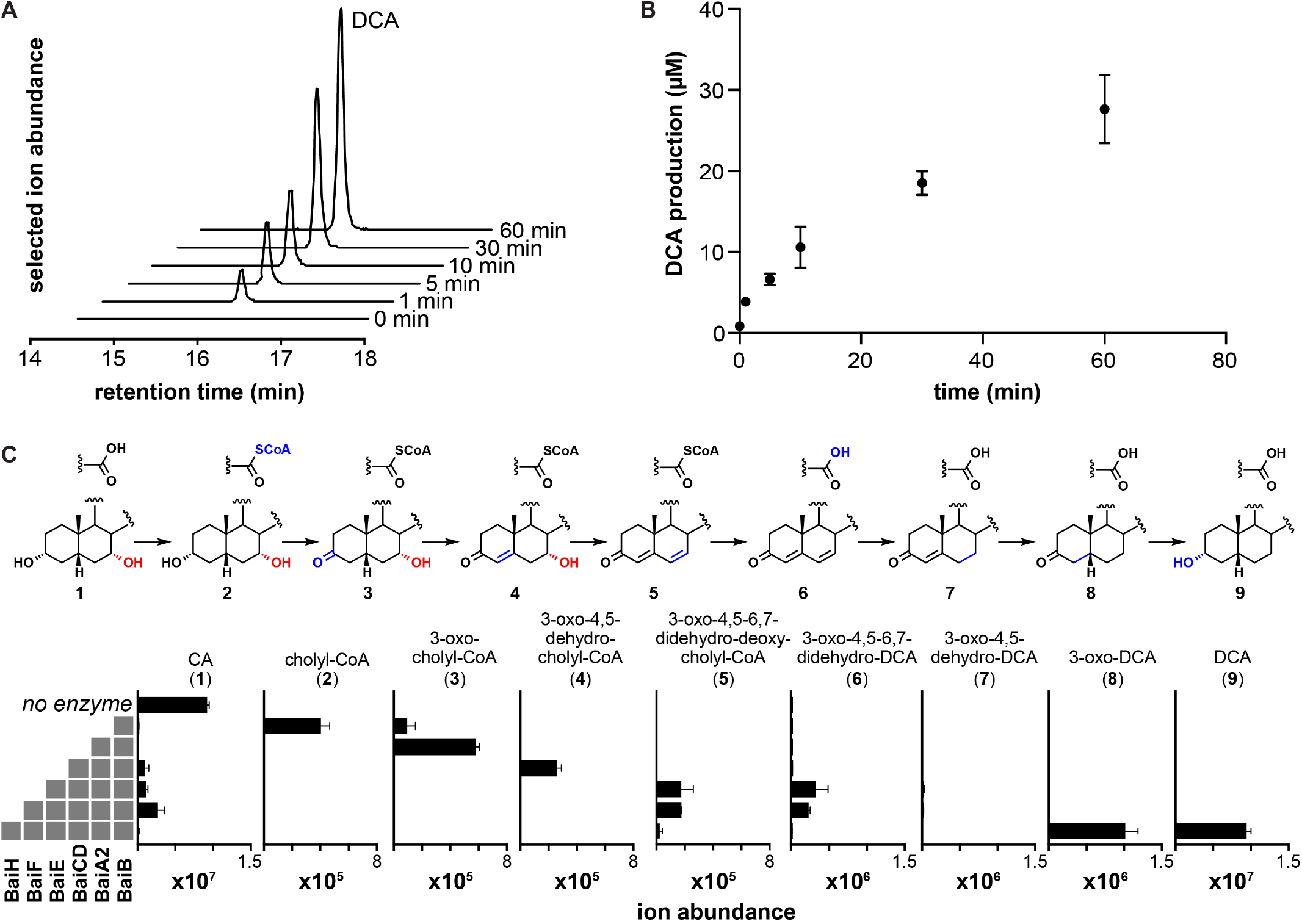
Establishing the complete 7α-dehydroxylation pathway in vitro. (A) EICs showing time-dependent production of DCA by six purified Bai enzymes. BaiB, BaiCD, BaiA2, BaiE, BaiF, and BaiH were purified and assayed anaerobically in the presence of NAD+, CoA, and ATP. Reactions were initiated by the addition of CA, and aliquots were analyzed by LC-MS at the indicated timepoints. (B) Time course of DCA production by a mixture of BaiB, BaiCD, BaiA2, BaiE, BaiF, and BaiH. Data points indicate the average level of DCA ± one SD (three biological replicates). (C) LC-MS ion abundance for DCA and pathway intermediates produced by a step-wise reconstitution assay in which the indicated enzymes were co-incubated as described in (A). Error bars indicate mean ± SD of three replicates.

To test our hypotheses regarding the order of steps in the pathway, we performed stepwise reconstitutions in which enzymes were added one at a time and intermediates were allowed to build up at each step in the pathway (**Fig. 2c**). From these data, we draw two conclusions: First, the six enzymes used in the reconstitution are not just sufficient but also necessary, and the pathway proceeds according to the scheme shown in **Fig. 4**. We directly observed mass ions consistent with each of the proposed intermediates, providing direct evidence for the previously proposed portion of the biosynthetic route. In spite of its conservation in all known dehydroxylating species, BaiI is dispensable for CA dehydroxylation *in vitro*. Since BaiI is a predicted Δ^5^-ketosteroid isomerase, it may process a substrate other than CA, likely one with a 4,5-or 5,6-olefin.

Second, to our surprise, the absence of BaiH caused the pathway to stall at the highly oxidized intermediate 3-oxo-4,5-6,7-didehydro-DCA, and its addition resulted in two successive 2e-reductions to form 3-oxo-DCA. BaiH had previously been proposed to oxidize an alternative substrate, 3-oxo-4,5-dehydro-UDCA^25^, so a potential role in the reductive arm of the pathway was unexpected. To explore this finding further, we incubated purified BaiH with synthetic 3-oxo-4,5-6,7-didehydro-DCA; we observed that the enzyme catalyzes a 2e-reduction to 3-oxo-4,5-dehydro-DCA, but does not reduce this intermediate further (**Fig. 3e**). Notably, 3-oxo-4,5-dehydro-DCA does not build up in the reconstitution reaction containing BaiH, suggesting that another enzyme present in the mixture catalyzes the second reductive step. Hypothesizing that the BaiH homolog BaiCD catalyzes the second reductive step, we incubated it with synthetic 3-oxo-4,5-dehydro-DCA, revealing that it reduces this substrate to 3-oxo-DCA (**Extended Data Fig. 3**). Together, these data show that the pathway employs an unusual redox strategy in which the A and B rings of the steroid core are converted into a highly oxidized intermediate, 3-oxo-4,5-6,7-didehydro-DCA; and that the two key reductive steps are catalyzed by two homologous enzymes in the Fe-S flavoenzyme superfamily, BaiH and BaiCD.

Finally, the last step in the pathway—reduction of 3-oxo-DCA to DCA—is carried out by BaiA2, as confirmed by assaying purified BaiA2 alone (**Extended Data Fig. 4**). Thus, BaiA2 and BaiCD both act twice in the pathway, catalyzing its first two and last two redox steps.

### Engineering the 7α-dehydroxylation pathway into *C. sporogenes*

Having determined the set of enzymes that are necessary and sufficient for the pathway, we sought to gain genetic control over the pathway as a first step toward engineering the bile acid output of the gut community. We began by attempting to construct a mutation in the *baiCD* gene of the native producer, *C. scindens*, using the ClosTron group II intron system; however, we were unsuccessful due to an inability to introduce DNA constructs into *C. scindens* by conjugation. As an alternative approach, we considered expressing the *bai* pathway in a gut commensal that is unable to carry out 7α-dehydroxylation; however, methods for transferring pathways in *Clostridium* are underdeveloped. The pathways for isobutanol (five genes) and 1,3-propanediol (three genes) have been transferred into *C. cellulolyticum* and *C. acetobutylicum*^28,29^, and a functional mini-cellulosome operon was introduced into the genome of *C. acetobutylicum*^30^. But to our knowledge, no more than a few genes have been transferred into *Clostridium*, and no pathway from the human microbiome has been mobilized from one *Clostridium* species to another.

We selected *Clostridium sporogenes* ATCC 15579 as the recipient for two reasons: it is related to *C. scindens*, making it likely that ancillary metabolic requirements for the pathway (e.g., cofactor biogenesis) would be met; and genetic tools have been developed that enable plasmids to be transformed into *C. sporogenes*^31^. Our initial attempts to clone the entire 8-gene *bai* operon (*baiB-baiI*) into an *E. coli-C. sporogenes* shuttle vector failed to yield clones harboring the complete operon. Reasoning that there might be a gene in the cluster that is toxic to *E. coli*, we cloned various fragments of the cluster under the control of different promoters (detailed in **Supplementary Table 1**), eventually managing to split the cluster into three pieces, each in its own *E. coli-C. sporogenes* shuttle vector: *baiB-baiF* in pMTL83153 (pMF01), *baiG* in pMTL83353 (pMF02), and *baiH-baiI* in pMTL83253 (pMF03) (**Fig. 3a**, **Extended Data Fig. 5**). Genes in pMF01 and pMF03 were placed under the control of the *spoIIE* promoter from *C. sporogenes* ATCC 15579, which is expressed during the late stages of *Clostridium* growth^32^, while *baiG* in pMF02 was driven by the strong *fdx* promoter. We conjugated these plasmids sequentially into *C. sporogenes* to yield strain MF001.

**Fig. 3.**
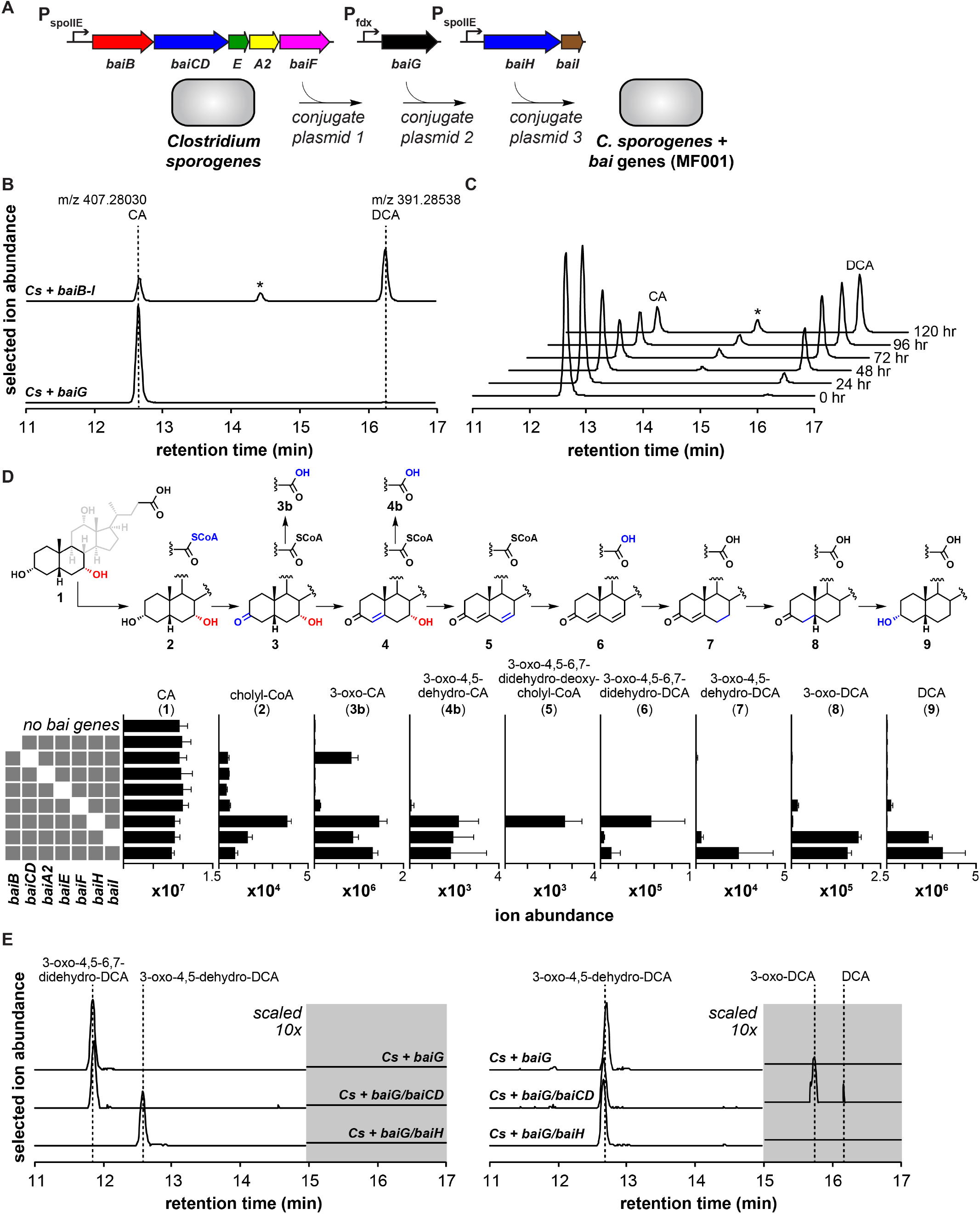
Transferring the 7α-dehydroxylation pathway in *Clostridium sporogenes*. (A) The *bai* operon was divided among three plasmids: *baiB-baiF* in pMTL83153 (pMF01), *baiG* in pMTL83353 (pMF02), and *baiH-bail* in pMTL83253 (pMF03). pMF01, pMF02, and pMF03 were conjugated successively into *Clostridium sporogenes* ATCC 15579 to create MF001. (B) Combined extracted ion chromatograms (EICs) showing the conversion of CA to DCA by MF001 versus a control strain of *C. sporogenes* harboring the transporter *baiG* (MF012). The strains were grown with 1 μM CA for 72 hr, extracted with acetone, and analyzed by LC-MS. The asterisk indicates isoDCA. (C) Combined EICs showing time-dependent conversion of CA to DCA by MF001. The strain was grown with 1 μM CA and aliquots from the indicated timepoints were analyzed as described in (B). (D) LC-MS ion abundances are shown for DCA, pathway intermediates, and derivatives produced by *C. sporogenes* strains with single gene deletions within the *bai* operon. Error bars indicate mean ± SD of three replicates. (E) Combined EICs showing the conversion of 3-oxo-4,5-6,7-didehydro-DCA to 3-oxo-4,5-dehydro-DCA by *C. sporogenes* + *baiG/baiH* (left), and the conversion of 3-oxo-4,5-dehydro-DCA to 3-oxo-DCA by *C. sporogenes* + *baiG/baiCD* (right). Each strain was cultivated with synthetic 3-oxo-4,5,6,7-didehydro-DCA (left) or 3-oxo-4,5-dehydro-DCA (right) for 72 hr and culture extracts were analyzed as in (B)

When incubated with CA, MF001 produces DCA in a time-dependent manner, in contrast to a control strain that harbors only the transporter (*baiG*) (**Fig. 3b, c**), which does not. Additionally, MF001 converts CDCA to LCA (**Extended Data Fig. 6**). These data show that the eight genes in the core *bai* cluster (**Fig. 1**) are sufficient to confer bile acid 7α-dehydroxylation on *C. sporogenes*, although they do not rule out the participation of one or more genes endogenous to *C. sporogenes*.

### Identifying branch points in the 7α-dehydroxylation pathway

To uncover potential branch points for engineering the biosynthesis of non-native pathway products, we constructed a set of strains in which each of the eight genes were individually deleted (**Extended Data Fig. 5**). We grew these strains with CA and assayed their culture supernatant for the build-up of intermediates (**Fig. 3d**). Deletion of genes encoding enzymes in the oxidative arm of the pathway resulted in the buildup of early pathway intermediates, as expected. Two exceptions were the *baiE* mutant, which produced only cholyl-CoA, potentially due to a polar effect on the transcription of downstream genes; and the baiF-deficient strain, which generated a small quantity of the final product DCA, suggesting there might be a compensatory CoA hydrolase (of note, *C. sporogenes* harbors two *baiF* homologs, CLOSPO_02756 and CLOSPO_00308), or that non-enzymatic hydrolysis of the CoA thioester happens to some extent *in vivo*.

Intriguingly, the *baiH* mutant accumulates a key intermediate in the reductive arm of the pathway, 3-oxo-4,5-6,7-didehydro-DCA (**Fig. 3d**), supporting our finding that BaiH catalyzes the first reductive step in the pathway and providing genetic access to a key intermediate in the pathway. Moreover, strains of *C. sporogenes* expressing BaiG/BaiH and BaiG/BaiCD convert, respectively, 3-oxo-4,5-6,7-didehydro-DCA to 3-oxo-4,5-dehydro-DCA and 3-oxo-4,5-dehydro-DCA to 3-oxo-DCA (**Fig. 3e**), providing access to intermediates that do not accumulate in a culture of *C. scindens*. Notably, the fully oxidized and partially reduced intermediates are branch points for the production of alternative bile acid metabolites including *allo* (5α) bile acids, which have important biological activities including the induction of regulatory T cells^33^. Thus, gaining genetic control over the pathway by expressing it in an alternative gut provides opportunities for rational and deliberate control of bile acid metabolism and the production of alternative molecules with distinct biological properties.

### 7α-dehydroxylation as a model for other microbiome-derived pathways

These results establish the complete 7α-dehydroxylation pathway, bringing this pathway closer to the level of knowledge we have about endogenous human metabolic pathways. Our work underscores that little is known about the biochemistry of metabolic pathways in the microbiome, in spite of the fact that they operate inside the human organism and produce abundant molecules that modulate host biology.

Key features of the bile acid 7α-dehydroxylation pathway might serve as a model for other pathways that produce high-abundance metabolites in the gut. We demonstrate that the reductive half of the pathway, which was previously uncharacterized, is centered around two reductions catalyzed by members of the Fe-S flavoenzyme superfamily^34^. Importantly, Fe-S flavoenzymes are known for shuttling electrons from the membrane to an organic (non-O2) terminal electron acceptor, enabling an anaerobic electron transport chain^35^. Moreover, the chemical logic of 7α-dehydroxylation is similar to that of other pathways used to support anaerobic electron transport chains. Here, a 4e-oxidation along the bottom of the A and B rings creates an enone with an acidic γ-proton, setting up a vinylogous dehydration of the 7-hydroxyl. The resulting dienone undergoes a 6e-reduction (three successive 2e-reductions), which nets the organism a 2e-reduction per molecule of primary bile acid. The key intermediate—an α,β-γ,δ-unsaturated ketone—is chemically similar to other oxidized intermediates that serve as electron acceptors in pathways from the microbiome, including fumarate, the electron acceptor for the *Bacteroides* anaerobic electron transport chain^36^; and aryl acrylic acids, which are electron acceptors for a subset of anaerobic Firmicutes^35^ (**Fig. 4**). The extended conjugation in these α,β-and α,β-γ,δ-unsaturated molecules allows them to serve as efficient electron acceptors. Although their redox potentials are lower than O2, they are well-suited for an anaerobic niche in which diffusible organic molecules are the most readily accessible alternative.

**Fig. 4.**
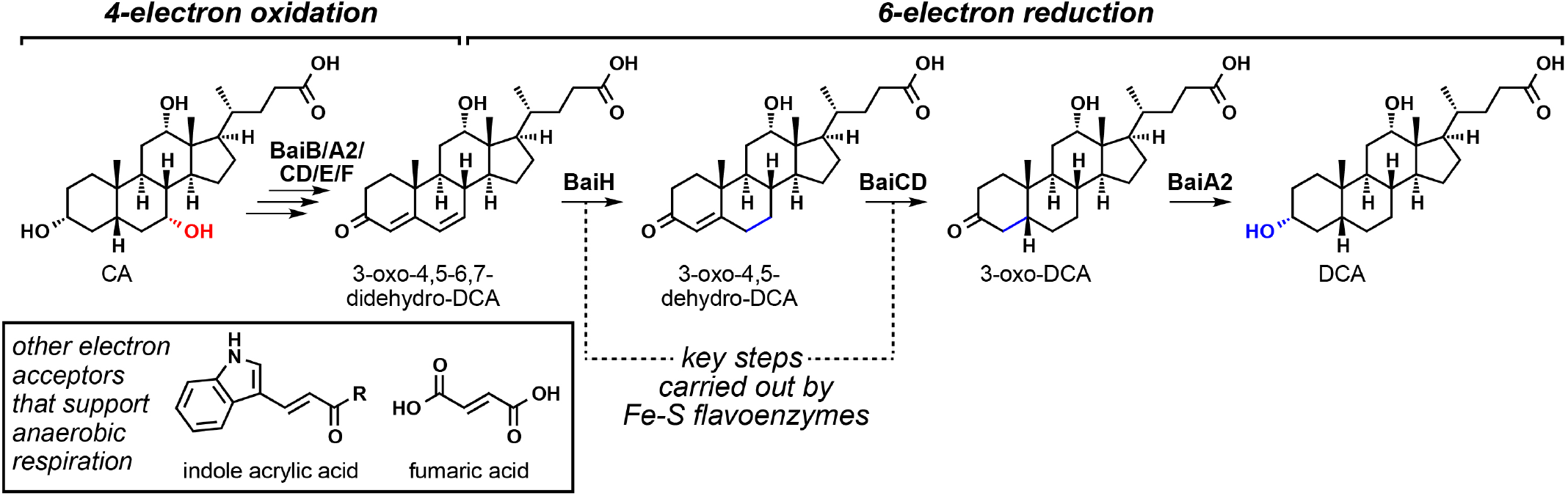
Highly oxidized metabolic intermediates as anaerobic electron acceptors. In the first half of the 7α-dehydroxylation pathway, two successive two-electron oxidations set up a vinylogous dehydration of the 7-hydroxyl, yielding the highly oxidized intermediate 3-oxo-4,5-6,7-didehydro-DCA. In the second half of the pathway, three successive two-electron reductions reduce this molecule to DCA, resulting in a net 2e-reduction. The first two of these reductions are carried out by Fe-S flavoenzymes, which harbor a suite of four cofactors that enable them to convert two-electron inputs to a one-electron manifold. A more detailed version of the previously proposed pathway is shown in **Extended Data Fig. 1**

### Engineering pathways from the microbiome

The gut microbiome harbors hundreds of pathways, many of which likely modulate host biology, but to date these pathways have not been a target of engineering. This stands in contrast to natural product pathways from terrestrial and marine microorganisms and plants, which are commonly expressed in heterologous hosts^37,38^ and engineered to generate non-native products^39^. Two technology gaps need to be overcome in order to make microbiome-derived pathways amenable to engineering: (i) efficient strategies for identifying pathways for known metabolites and small molecule products of orphan gene clusters, and (ii) tools for transferring pathways into bacterial hosts native to the gut and manipulating them to produce novel molecules. The work described here is a starting point for these efforts, and provides set of tools for deorphaning, heterologously expressing, and engineering pathways from *Clostridium* as a new way of controlling the chemical output of the microbiome.

## METHODS

### Bacterial strains, culture conditions, and bile acids

*Clostridium scindens* VPI 12708 and *Clostridium sporogenes* ATCC 15579 were obtained from the Japan Collection of Microorganisms (JCM) and the American Type Culture Collection (ATCC), respectively. Engineered *C. sporogenes* strains used in this study are shown in **Supplementary Table 2**. They were cultured in TYG (3% w/v tryptone, 2% w/v yeast extract, 0.1% w/v sodium thioglycolate) broth at 37 °C in an anaerobic chamber from Coy Laboratories. *Escherichia coli* CA434 (HB101/pRK24) was cultured at 37 °C in LB broth supplemented with 12 μg/mL tetracycline and 100 μg/mL carbenicillin. In addition, 20 μg/mL chloramphenicol, 100 μg/mL spectinomycin or 250 μg/mL erythromycin was used for the selection of series of plasmids of pMTL83153, pMTL83353 or pMTL83253 respectively. Plasmids used in this study are shown in **Supplementary Table 1**. Cholic acid (1), chenodeoxycholic acid, deoxycholic acid (9) and lithocholic acid were purchased from Sigma-Aldrich. 3-oxo-cholic acid (3b) and 3-oxo-deoxycholic acid (8) were purchased from Steraloids. 3-oxo-4,5-6,7-didehydro-DCA (6) and 3-oxo-4,5-dehydro-DCA (7) were synthesized using previously reported procedures^40^. Structural assignments for the remaining pathway intermediates and derivatives shown in **Fig. 2** and **Fig. 3** are provisional, and were made on the basis of mass spectra, retention times, and comparison to chemically related standards.

### Cloning of the *bai* operon

All PCR amplification was conducted using PrimeSTAR Max DNA polymerase (Takara Bio) according to the manufacturer’s instructions. Sequences of primers for target genes and cloning vectors were shown in **Supplementary Table 3**. For the heterologous expression of *bai* genes under fdx promoter, pMTL vectors were amplified with primers 1 and 2. For the expression of *bai* genes under spoIIE promoter, pMTL vectors harboring spoIIE promoter was constructed at first. pMTL vectors were amplified with primers 1 and 3 to remove the fdx promoter and spoIIE promoter region, which is the 277 bp sequence upstream of CLOSPO_01065, was amplified with primers 4 and 5. Then these two PCR fragments were assembled by overlap PCR. The target gene sequences were amplified with the primers pair shown in **Supplementary Table 3**. PCR fragments were assembled with the amplified fragments of vectors using Gibson assembly kit (New England Bio Labs). *E. coli* Stbl4 competent cells (Invitrogen) were transformed with the assembled plasmids by electroporation and transformants were confirmed by PCR. Positive clones harboring assembled plasmids were cultivated, and the plasmid was obtained by miniprep and verified by sequencing.

### Heterologous expression in *C. sporogenes*

For the heterologous expression experiments, plasmids were transferred into *C. sporogenes* by conjugation using *E. coli* CA434. *E. coli* CA434 was electroporated with the individual plasmids and recovered overnight in selective media. 1 mL of overnight culture of the resultant transformants was harvested. The cell pellet was washed with PBS to remove residual antibiotics and re-suspended with 200 μL of an overnight culture of *C. sporogenes* in anaerobic chamber. Eight drops of 25 μL of the suspension were pipetted on TYG agar plate without antibiotics and the plate was incubated anaerobically at 37 °C for 2 days. The bacterial biomass was scraped up and resuspended in 300 μL of PBS. The whole cell suspension was then plated on TYG agar plates supplemented with 250 μg/mL D-cycloserine and appropriate antibiotics (15 μg/mL thiamphenicol for pMTL83153, 500 μg/mL spectinomycin for pMTL83353 or 5 μg/mL erythromycin for pMTL83253). After a few days, antibiotic resistant colonies were picked and re-streaked on agar containing the same antibiotic. The resulting clones were confirmed by PCR amplification using appropriate primers (**Supplementary Table 3**). Multiple plasmids were introduced sequentially, using the same procedure.

### Extraction of metabolites

Engineered strains were cultured anaerobically in TYG medium supplemented with appropriate antibiotics from frozen glycerol stocks. 10 μL of the overnight culture was inoculated in 1 mL of TYG medium supplemented with appropriate antibiotics and 1 μM substrate. After 72 hr, unless otherwise noted, the culture was extracted with 20% acetone and centrifuged. The supernatant was analyzed by LC/MS.

### LC/MS analysis of metabolite extracts

Metabolite extracts were analyzed using an Agilent 1290 LC system coupled to an Agilent 6530 QTOF with a 1.7 μm, 2.1 × 100 mm Kinetex C18 column (Phenomenex). Water with 0.05% formic acid (A) and acetone with 0.05% formic acid (B) was used as the mobile phase at a flow rate of 0.35 mL/min over a 32 min gradient: 0-1 min, 25% B; 1-25 min, 25-75% B; 25-26 min, 75-100% B; 26-30 min, 100% B; 30-32 min 75-25% B. All data were collected in negative ion mode.

For detection of CoA conjugates, a 1.8 μm, 2.1 × 50 mm ZORBAX SB-C18 column (Agilent Technologies) and water with 10 mM ammonium acetate pH 9.0 (A) and acetonitrile (B) was used. A flow rate of 0.3 mL/min was used over the 17 min gradient: 0-2 min, 15% B; 2-14 min, 15-50% B; 14-14.1 min 50-95% B, 14.1-17 min, 85% B. All data were collected in positive ion mode.

### Cloning of *bai* operon genes

To increase the probability of assembling a complete bai operon, we cloned the genes encoding *baiB, baiA2, baiCD, baiE, baiF*, and *baiH* from *Clostridium scindens* VPI12708, *Clostridium hylemonae*, and *Clostridium hiranonis* using the primers in **Supplementary Table 4** and the KOD Xtreme^™^ Hot Start PCR kit (Millipore) following the manufacturers protocol. Each PCR amplified gene contains ligation independent cloning (LIC) sites that are complimentary to the pSGC vector. The PCR products were purified with the Agencourt Ampure XP PCR clean-up kit (Beckman Coulter) according to the manufacturers protocol. The pSGC vector was prepared for LIC by linearization with the restriction enzyme BsaI. LIC sites were installed by adding T4 DNA polymerase (NEB) to 10 μg of linearized plasmid in a 50 μL reaction containing 2.5 mM GTP, 1 × NEB Buffer 2, and 1 × BSA for 1 hr at 22 °C. T4 DNA polymerase was heat-inactivated by incubation at 75 °C for 20 min. The 2 μL of the PCR products were also treated with T4 DNA polymerase in a 10 μL reaction containing 2.5 mM CTP, 1 × NEB Buffer 2, and 1 × BSA for 1 hr at 22 ° C. T4 DNA polymerase was heat inactivated. The LIC reaction was carried out by mixing 15 ng of digested vector with ~ 40 ng of digested PCR product with a subsequent incubation at 22 °C for 10 min. A 30 μL aliquot of DH10b cells (NEB) were transformed with 2 μL of the above mixture using standard bacterial transformation protocols. Cloning the genes into pSGC with this method adds a His6 tag to the N-terminus of each protein with the following sequence: MHHHHHHSSGVDLGTENLYFQS. All final constructs were sequence-verified (Genescript).

### Expression and purification of BaiH and BaiCD

BL-21(DE3) cells containing the pPH151 plasmid were transformed with the pSGC plasmid containing either BaiCD or BaiH. The transformants were selected on an LB/agar plate containing 50 μg/mL kanamycin and 34 μg/mL chloramphenicol. A single colony was used to inoculate 20 mL of LB overnight culture containing the above antibiotics. The overnight culture was used to inoculate 2 L of Studier’s auto induction media (ZYP-5052 supplemented with 1 mM flavin mononucleotide and 200 μM FeCl3) housed in a 2 L PYREX® media bottle. Cultures were grown with constant aeration using a sparging stone attached to a pressurized, 0.22 μm filtered air source all in a water bath maintained at 37 °C. After 5 hr, aeration was stopped and the culture was placed in an ice bath for 1 hr. The culture was returned to a 22 °C water bath and light aeration was resumed. After 5 min, cysteine was added to a final concentration of 600 μM. The culture was grown at 22 °C for ~ 20 hr before being harvested by centrifugation at 10,000 × g. Cell pellets were flash frozen and stored in liquid N2 until purification. All subsequent steps were carried out in an MBraun anaerobic chamber maintained at < 0.1 ppm oxygen (MBraun, Stratham, NH). Plastics were brought into the chamber and allowed to sit for two weeks before use. All solvents and buffer stocks were degassed by sparging with argon gas for 4 hr before being taken into the chamber. In a typical purification, ~30 grams of BaiCD or BaiH cell paste was resuspended in 30 mL of lysis buffer containing 50 mM HEPES, pH 7.5, 300 mM KCl, 4 mM imidazole, 10 mM 2-mercaptoethanol (BME), 10% glycerol, 1 mM FMN, 1 mM FAD, and 1% Triton-X305. The resuspension was subjected to 50 rounds of sonic disruption (80% output, 3 s pulse on, 12 s pulse of) at 4 °C. The lysate was cleared by centrifugation at 4 °C for 1 hr at 15,000 × g. The supernatant was loaded with an ÄKTA express FPLC system onto a 5 mL fast-flow HisTrap^™^ column (GE Healthcare Life Sciences) equilibrated in lysis buffer lacking FMA, FAD, and Triton-X305. The column was washed with 10 column volumes of lysis buffer before elution with 5 mL of buffer containing 50 mM HEPES, pH 7.5, 300 mM KCl, 300 mM imidazole, 10 mM BME, and 10% glycerol. The fractions containing protein, based on absorbance at 280 nm, were pooled and reconstituted with Fe and sulfur as previously described. The reconstituted proteins were then passed over a HiPrep 16/60 Sephacryl S-200 HR column equilibrated in 20 mM HEPES, pH 7.5, 300 mM KCl, 5 mM DTT, and 10% glycerol. The proteins were concentrated to ~ 1 mL with a vivaspin 20 concentrator (Sartorius Stedium Biotech). The protein concentration was estimated by A280 using the extinction coefficient calculated based on its corresponding amino acid sequence.

### Expression and purification of BaiB, BaiA2, BaiE, and BaiF

BL-21(DE3) cells containing the pRIL plasmid were transformed with the plasmid containing BaiB, BaiA2, BaiE, or BaiF. Each transformant was selected on an LB/agar plate containing 50 μg/mL kanamycin and 34 μg/mL chloramphenicol. A single colony was used to inoculate 20 mL of LB overnight culture containing the above antibiotics. The overnight culture was used to inoculate 2 L of Studier’s auto induction media (ZYP-5052) housed in a 2 L PYREX^®^ media bottle. Cultures were grown with constant aeration using a sparging stone attached to a pressurized, 0.22 μm filtered air source in a water bath at 37 °C. After 5 hr, aeration was stopped and the culture was placed in an ice bath for 1 hr. The culture was returned to a 22 °C water bath and light aeration was resumed. The culture was grown at 22 °C for ~ 20 hr before being harvested by centrifugation at 10,000 x g. Cell pellets were flash frozen and stored in liquid N2 until purification. All subsequent steps were carried out in an MBraun anaerobic chamber maintained at < 0.1 ppm oxygen as above with minor modifications. Briefly, a typical purification, ~ 30 – 40 grams of cell paste was resuspended in 30 – 40 mL of lysis buffer containing 50 mM HEPES, pH 7.5, 300 mM KCl, 4 mM imidazole, 10 mM 2-mercaptoethanol (BME), 10% glycerol, and 1% Triton-X305. The resuspension was subjected to 50 rounds of sonic disruption (80% output, 3 s pulse on, 12 s pulse of) at 4 °C. The lysate was cleared by centrifugation at 4 °C for 1 hr at 15,000 × g. The supernatant was loaded with an ÄKTA express FPLC system onto a 5 mL fast-flow HisTrap^™^ column (GE Healthcare Life Sciences) equilibrated in lysis buffer lacking Triton-X305. The column was washed with 10 column volumes of lysis buffer before elution with 5 mL of buffer containing 50 mM HEPES, pH 7.5, 300 mM KCl, 300 mM imidazole, 10 mM BME, and 10% glycerol. The fractions containing protein, based on absorbance at 280 nm, were pooled and immediately passed over a HiPrep 16/60 Sephacryl S-200 HR column equilibrated in 20 mM HEPES, pH 7.5, 300 mM KCl, 5 mM DTT, and 10% glycerol. The proteins were concentrated to ~ 1 mL with a vivaspin 20 concentrator (Sartorius Stedium Biotech). The protein concentration was estimated by A280 using the extinction coefficient calculated based on its corresponding amino acid sequence.

### Bile acid pathway *in vitro* reconstitution

Six assays each contained 50 mM HEPES pH 7.5, 50 mM KCl, 200 μM NAD, 100 μM CoA, and 200 μM ATP. In addition, each assay contained 0.1 mM of 1-6 of the following enzymes: BaiB from *C. scindens*, BaiA2 from *C. scindens*, BaiCD from *C. hiranonis*, BaiE from *C. hiranonis*, BaiF from *C. hylemonae*, and BaiH from *C. scindens*. All reactions were initiated with the addition of cholic acid and incubated at 22 °C for 30 min before being quenched by the additions of an equal volume of 100 % acetone. Each assay was performed in triplicate. Product formation was monitored by LC/MS described above.

### Bile acid pathway in vitro reconstitution kinetics

To determine the rate of DCA production by the *in vitro* pathway, assays were performed with 50 mM HEPES pH 7.5, 50 mM KCl, 200 μM NAD, 100 μM CoA, and 200 μM ATP, 0.1 mM of BaiB from *C. scindens*, BaiA2 from *C. scindens*, BaiCD from *C. hiranonis*, BaiE from *C. hiranonis*, BaiF from *C. hylemonae*, and BaiH from *C. scindens*. Reactions were initiated with the addition of cholic acid and incubated at 22 °C. Samples of the reaction were removed and mixed with an equal volume (100 mM H2SO4 at designated times. Each assay was performed in triplicate. Product formation was monitored by LC/MS described above.

### *K_M_* assay for BaiCD

Kinetic parameters for BaiCD from *C. hiranonis* were determined in assays that contained 1 μM enzyme, 50 mM HEPES pH 7.5, 50 mM KCl, and 500 μM NADH. Reactions mixtures were incubated for 5 min at 22 °C before being initiated with 3-oxo-4,5-dehydro-deoxycholic acid. Concentrations of substrate were varied between 3.91 μM and 500 μM. 20 μL samples were removed and mixed with an equal volume of 100 mM H2SO4 to stop the reaction. Product formation was determined by LC/MS described above. Reactions were performed in triplicate and the data were fit to the Michaelis-Menten equation by the least squares method.

### *K_M_* assay for BaiH

Kinetic parameters for BaiH from *C. scindens* were determined in assays that contained 0.45 μM enzyme, 50 mM HEPES pH 7.5, 50 mM KCl, and 500 μM NADH. Reactions mixtures were incubated for 5 min at 22 °C before being initiated with 3-oxo-4,5,6,7-didehydro-deoxycholic acid. Concentrations of substrate were varied between 0.78 μM and 100 μM. 20 μL samples were removed and mixed with an equal volume 100 mM H2SO4 to stop the reaction. Product formation was determined by LC/MS described above. Reactions were performed in triplicate and the data were fit to the Michaelis-Menten equation by the least squares method.

## SUPPLEMENTARY INFORMATION

Supplementary Tables 1-4 are available in the online version of the paper.

## ACKNOWLEDGMENTS

We are deeply indebted to Christopher T. Walsh, Dylan Dodd, Colleen O’Loughlin, and members of the Fischbach and Almo laboratories for helpful comments on the manuscript. This work was supported by NIH grants DP1 DK113598 (M.A.F.), R01 DK110174 (M.A.F.), P01 GM118303-01 (S.C.A.), U54 GM093342 (S.C.A.), and U54 GM094662 (S.C.A.), the Chan-Zuckerberg Biohub (M.A.F.), an HHMI-Simons Faculty Scholars Award (M.A.F.), an Investigators in the Pathogenesis of Infectious Disease award from the Burroughs Wellcome Foundation (M.A.F.), and the Price Family Foundation (S.C.A.).

## AUTHOR CONTRIBUTIONS

M.F., T.L.G., S.C.A., and M.A.F. conceived and designed the experiments. M.F., T.L.G., Y.V., M.M., L.C.B., and C.G. performed experiments. V.P., M.A.F., and M.H.M. conceived and designed the computational analyses. V.P. performed the computational analyses. M.F., T.L.G., V.P., M.H.M., S.C.A., and M.A.F. analyzed data and wrote the manuscript. All authors discussed the results and commented on the manuscript.

## AUTHOR INFORMATION

Reprints and permissions information is available at http://www.nature.com/reprints. The authors declare no competing financial interests. Correspondence and requests for materials should be addressed to S.C.A. or M.A.F.

---

**Extended Data Fig. 1.**
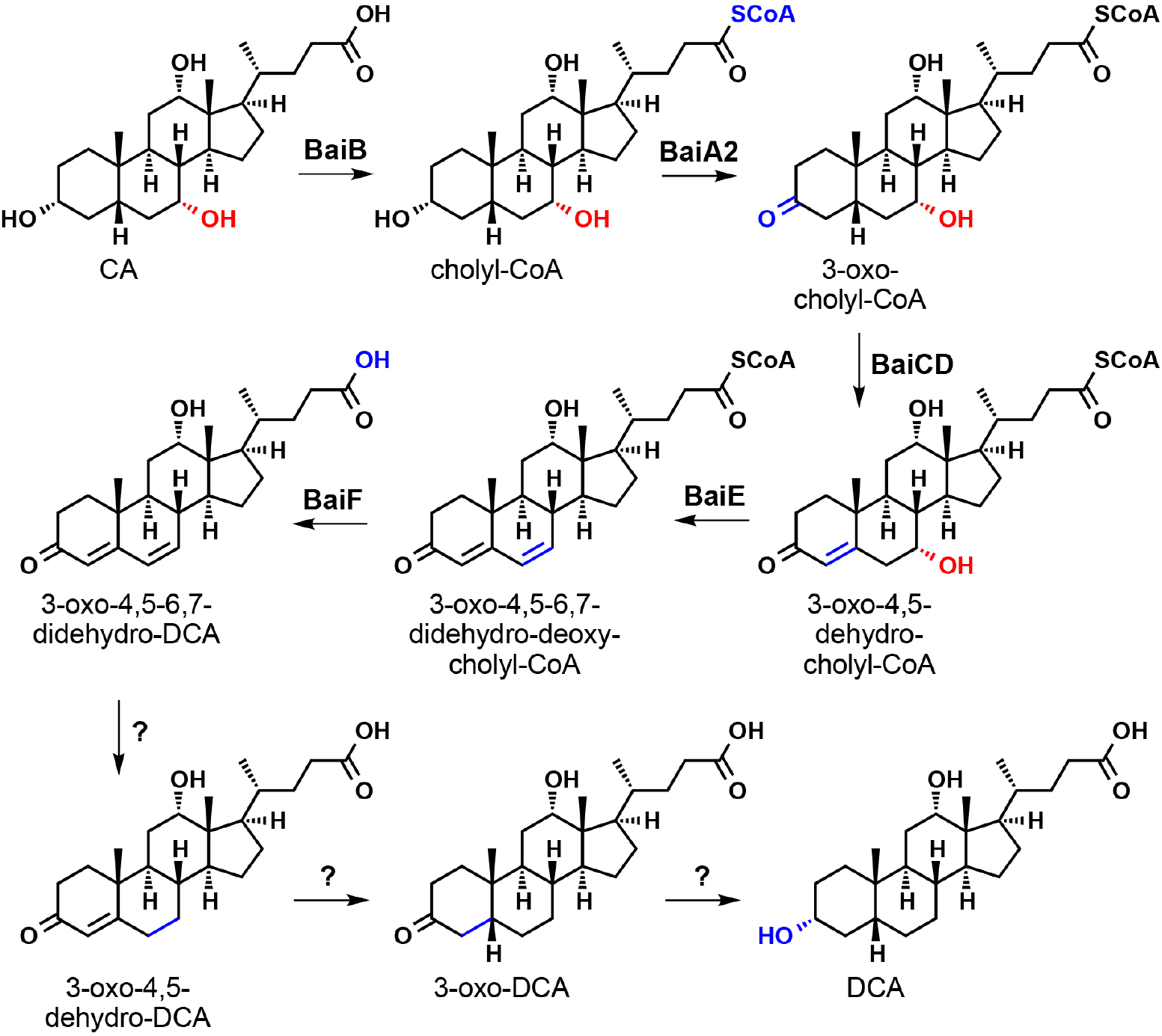
Previously proposed pathway for 7α-dehydroxylation of CA in *C. scindens* VPI 12708. See main text for details and a summary of the previous literature.

**Extended Data Fig. 2.**
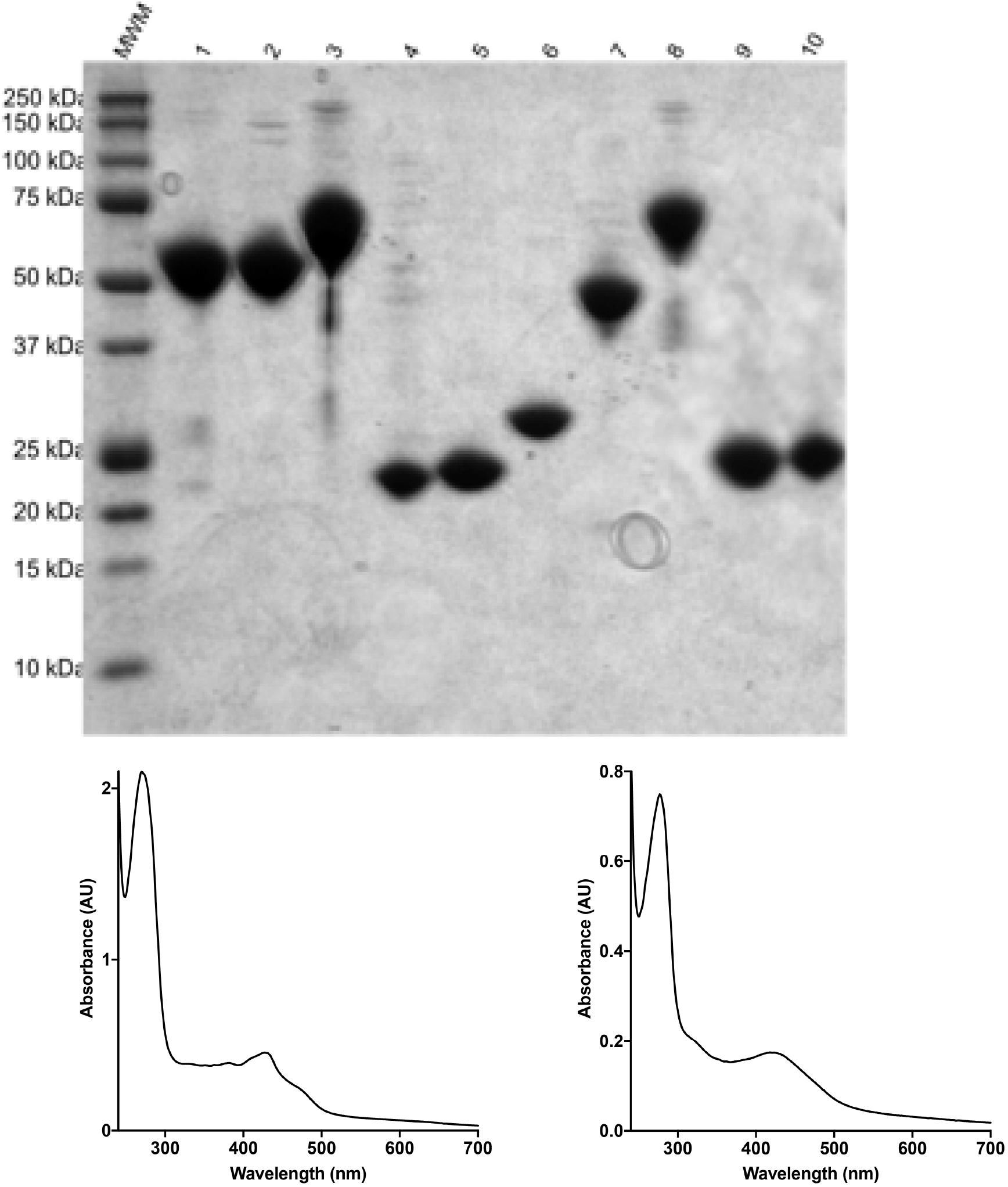
Purification of recombinant Bai proteins. (A) SDS-PAGE analysis of purified Bai proteins after Ni-affinity and size exclusion purification. MWM, molecular weight marker; 1, BaiB from *C. scindens;* 2, BaiB from *C. hylemonae;* 3, BaiCD from *C. hiranonis*; 4, BaiE from C. *scindens;* 5, BaiE from *C. hiranonis;* 6, BaiA2 from *C. scindens;* 7, BaiF from *C. hylemonae;* 8, BaiH from *C. scindens;* 9, BaiI from *C. scindens;* 10, BaiI from *C. hiranonis*. B) UV-visible spectra of BaiCD from *C. hiranonis* (left) and BaiH from *C. scindens* (right).

**Extended Data Fig. 3.**
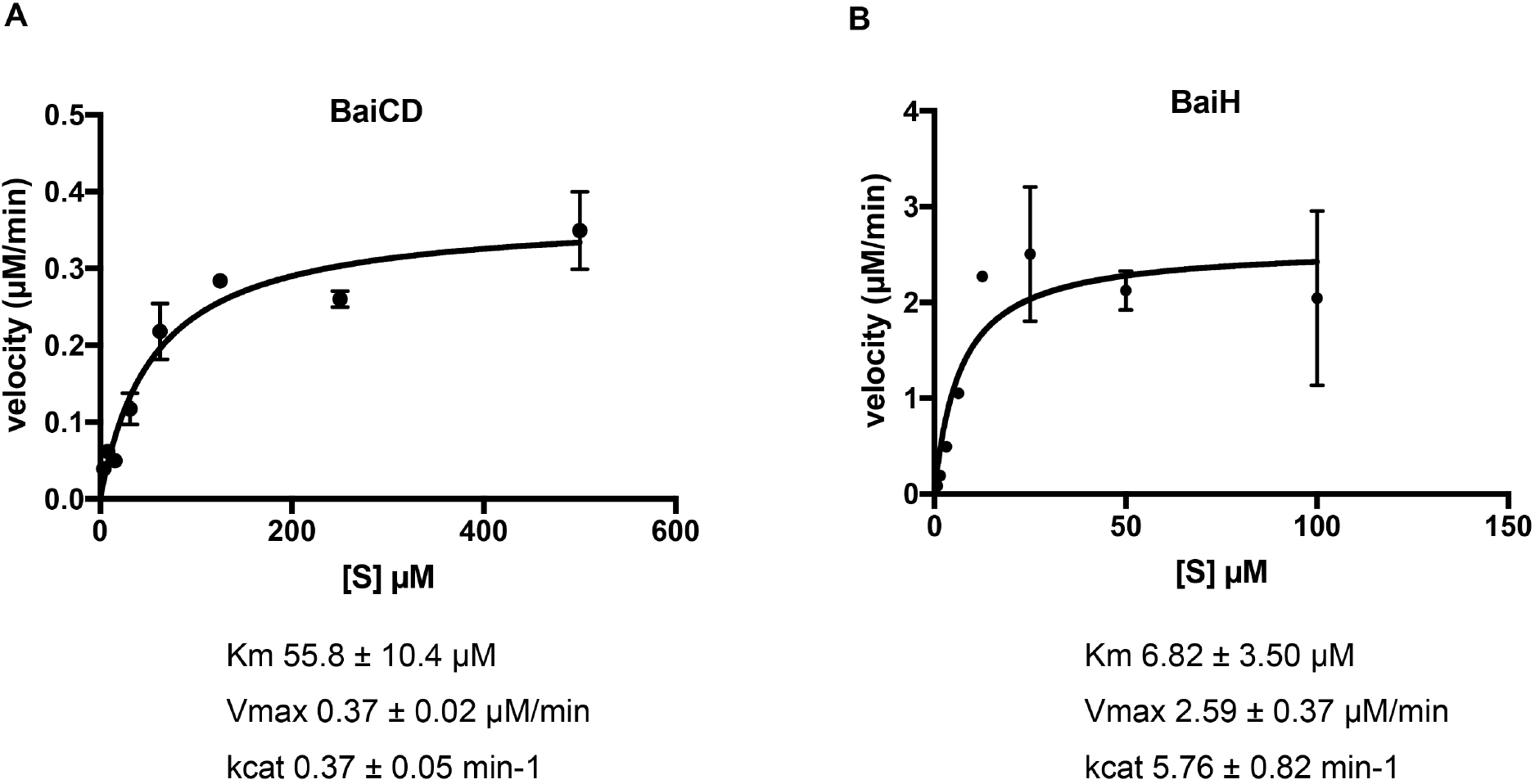
Kinetic parameters for BaiCD and BaiH. (A) Michaelis-Menten analysis of the conversion of 3-oxo-4,5-dehydro-DCA to 3-oxo-DCA by BaiCD. B) Michaelis-Menten analysis of the conversion of 3-oxo-4,5,6,7-didehydro-DCA to 3-oxo-4,5-dehydro-DCA by BaiH. Data indicate the average product level ± one SD (three biological replicates).

**Extended Data Fig. 4.**
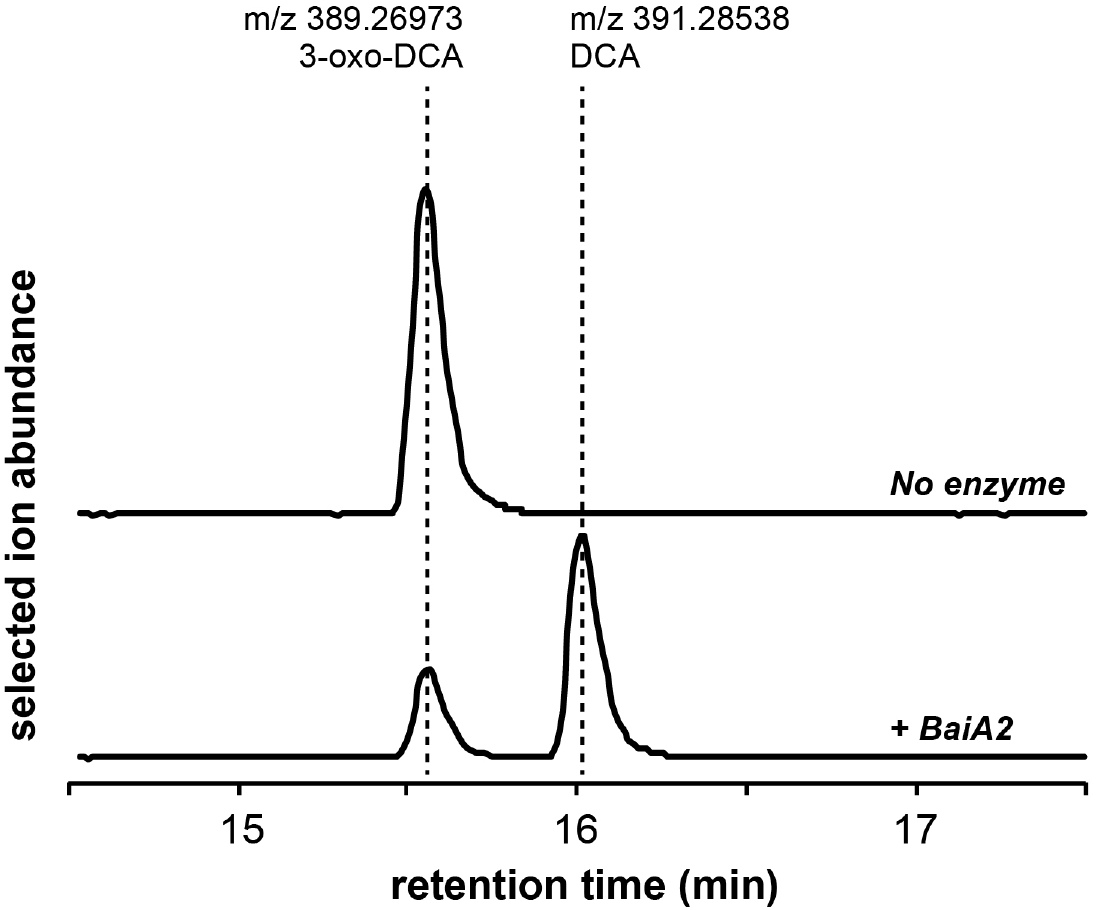
Biochemical analysis of 3-oxo-DCA reduction by BaiA2. Combined extracted ion chromatograms showing the conversion of 3-oxo-DCA to DCA by recombinant BaiA2.

**Extended Data Fig. 5.**
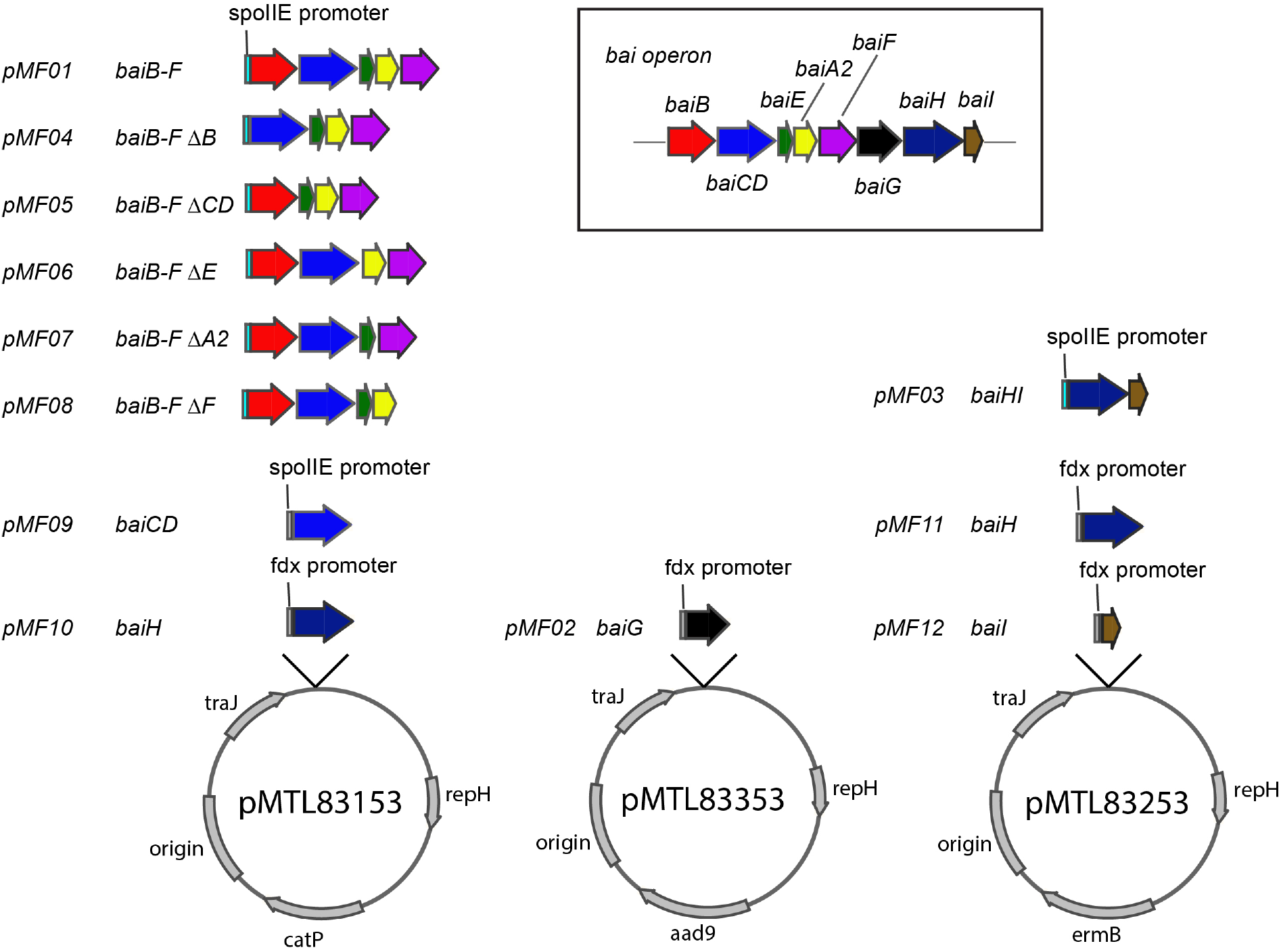
Constructs for expressing the *bai* operon and portions thereof in *C. sporogenes*. Each of the plasmids has replication origins for *E. coli* and *Clostridium*, traJ to enjable conjugal plasmid transfer, and an antibiotic resistance gene. *bai* genes were introduced into these plasmids under the control of the fdx or spoIIE promoter. For the genetic analysis of *baiCD* and *baiH* function, pMTL83153-based plasmids were used.

**Extended Data Fig. 6.**
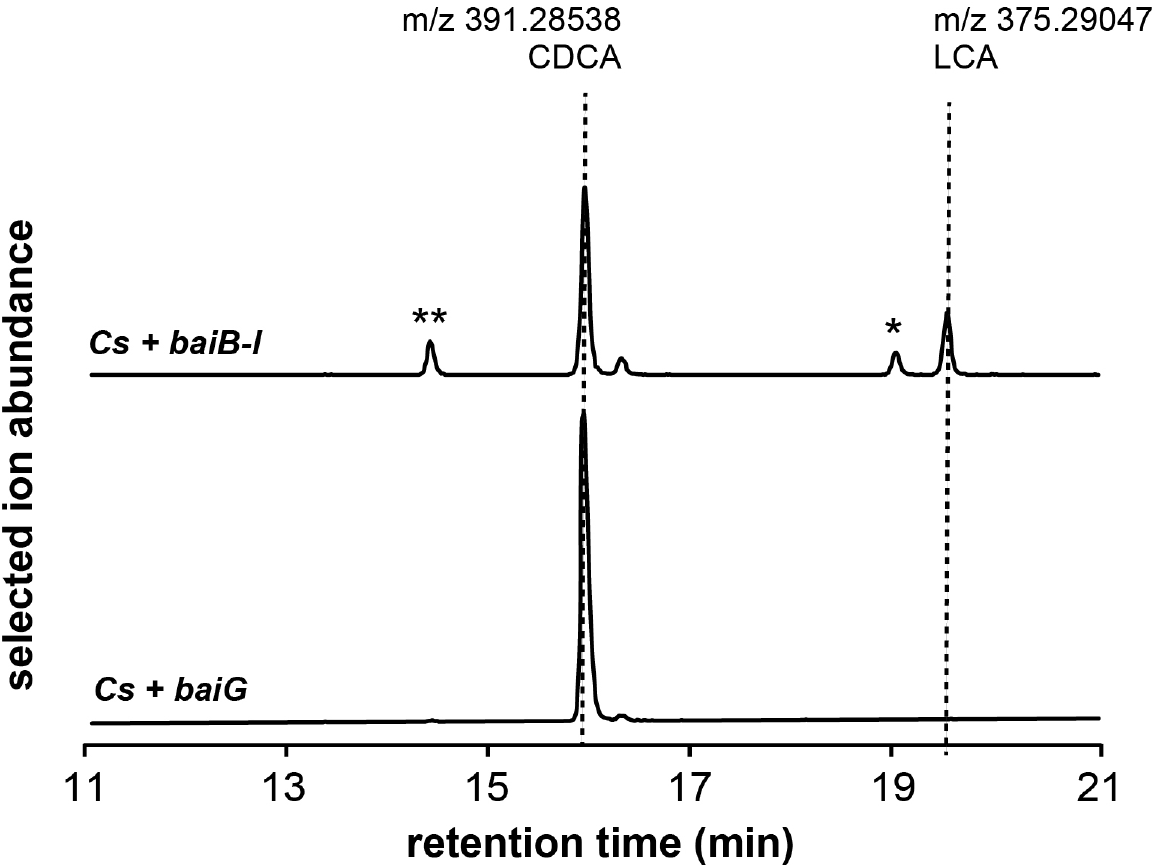
7α-dehydroxylation of CDCA *in vivo*. Combined extracted ion chromatograms showing the conversion of CDCA to LCA by a *C. sporogenes* strain harboring the complete *bai* operon on three plasmids (MF001) versus a control strain of *C. sporogenes* harboring the transporter *baiG* (MF012). The strains were cultivated with 1 μM CA for 72 hr; an acetone extract of the culture supernatant was analyzed by HPLC-MS. The single asterisk indicates isoLCA, and the double asterisk is provisionally assigned as isoCDCA.

**Supplementary Table 1.**

| Plasmid | Backbone | Promoter | Genes introduced | Source |
| --- | --- | --- | --- | --- |
| pMTL83153 | pMTL83153 | fdx |  | Nigel Minton |
| pMTL83353 | pMTL83353 | fdx |  | Nigel Minton |
| pMTL83253 | pMTL83253 | fdx |  | Nigel Minton |
| pMF01 | pMTL83153 | spolIE | *baiB_baiCD_baiE_baiA2_baiF* | This study |
| pMF02 | pMTL83353 | fdx | *baiG* | This study |
| pMF03 | pMTL83253 | spolIE | *baiH_bail* | This study |
| pMF04 | pMTL83153 | spolIE | *baiCD_baiE_baiA2_baiF* | This study |
| pMF05 | pMTL83153 | spolIE | *baiB_baiE_baiA2_baiF* | This study |
| pMF06 | pMTL83153 | spolIE | *baiB_baiCD_baiA2_baiF* | This study |
| pMF07 | pMTL83153 | spolIE | *baiB_baiCD_baiE_baiF* | This study |
| pMF08 | pMTL83153 | spolIE | *baiB_baiCD_baiE_baiA2* | This study |
| pMF09 | pMTL83153 | spolIE | *baiCD* | This study |
| pMF10 | pMTL83153 | fdx | *baiH* | This study |
| pMF11 | pMTL83253 | fdx | *baiH* | This study |
| pMF12 | pMTL83253 | fdx | *bail* | This study |

**Supplementary Table 2.**

| Strain | Plasmids | Genes |
| --- | --- | --- |
| MF001 | pMF01, pMF02, pMF03 | *baiB, baiCD, baiE, baiA2, baiF, baiG, baiH, bail* |
| MF002 | pMF04, pMF02, pMF03 | *baiCD, baiE, baiA2, baiF, baiG, baiH, bail* |
| MF003 | pMF05, pMF02, pMF03 | *baiB, baiE, baiA2, baiF, baiG, baiH, bail* |
| MF004 | pMF06, pMF02, pMF03 | *baiB, baiCD, baiA2, baiF, baiG, baiH, bail* |
| MF005 | pMF07, pMF02, pMF03 | *baiB, baiCD, baiE, baiF, baiG, baiH, bail* |
| MF006 | pMF08, pMF02, pMF03 | *baiB, baiCD, baiE, baiA2, baiG, baiH, bail* |
| MF007 | pMF01, pMF02, pMF12 | *baiB, baiCD, baiE, baiA2, baiF, baiG, bail* |
| MF008 | pMF01, pMF02, pMF11 | *baiB, baiCD, baiE, baiA2, baiF, baiG, baiH* |
| MF009 | pMF01, pMF02 | *baiB, baiCD, baiE, baiA2, baiF, baiG* |
| MF010 | pMF09, pMF02 | *baiCD, baiG* |
| MF011 | pMF10, pMF02 | *baiH, baiG* |
| MF012 | pMTL83153, pMF02, pMTL83253 | *baiG* |

**SI Table 3:**
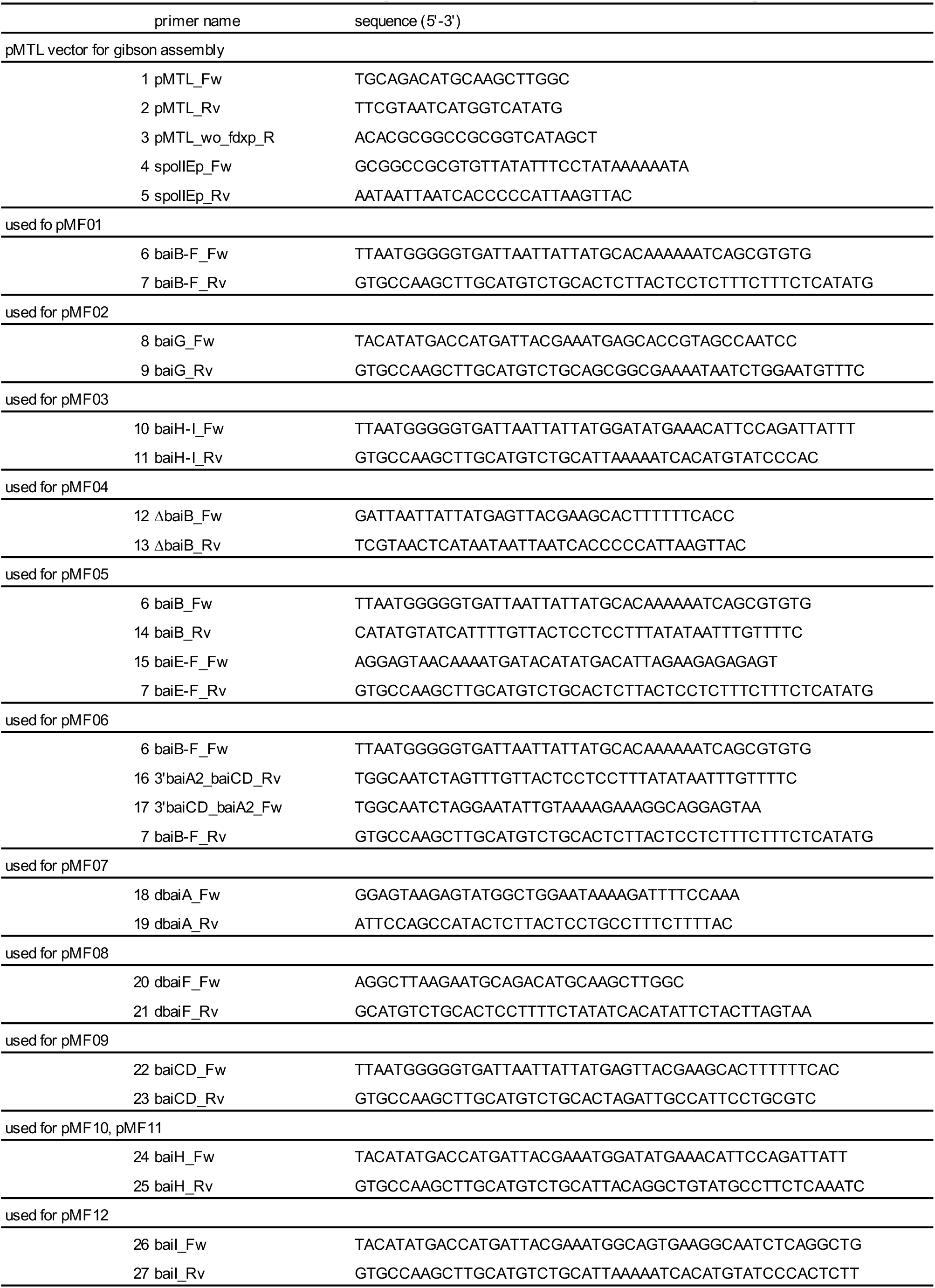
Primers used for heterologous expression vector for C. sporogenes

**SI Table 4:**
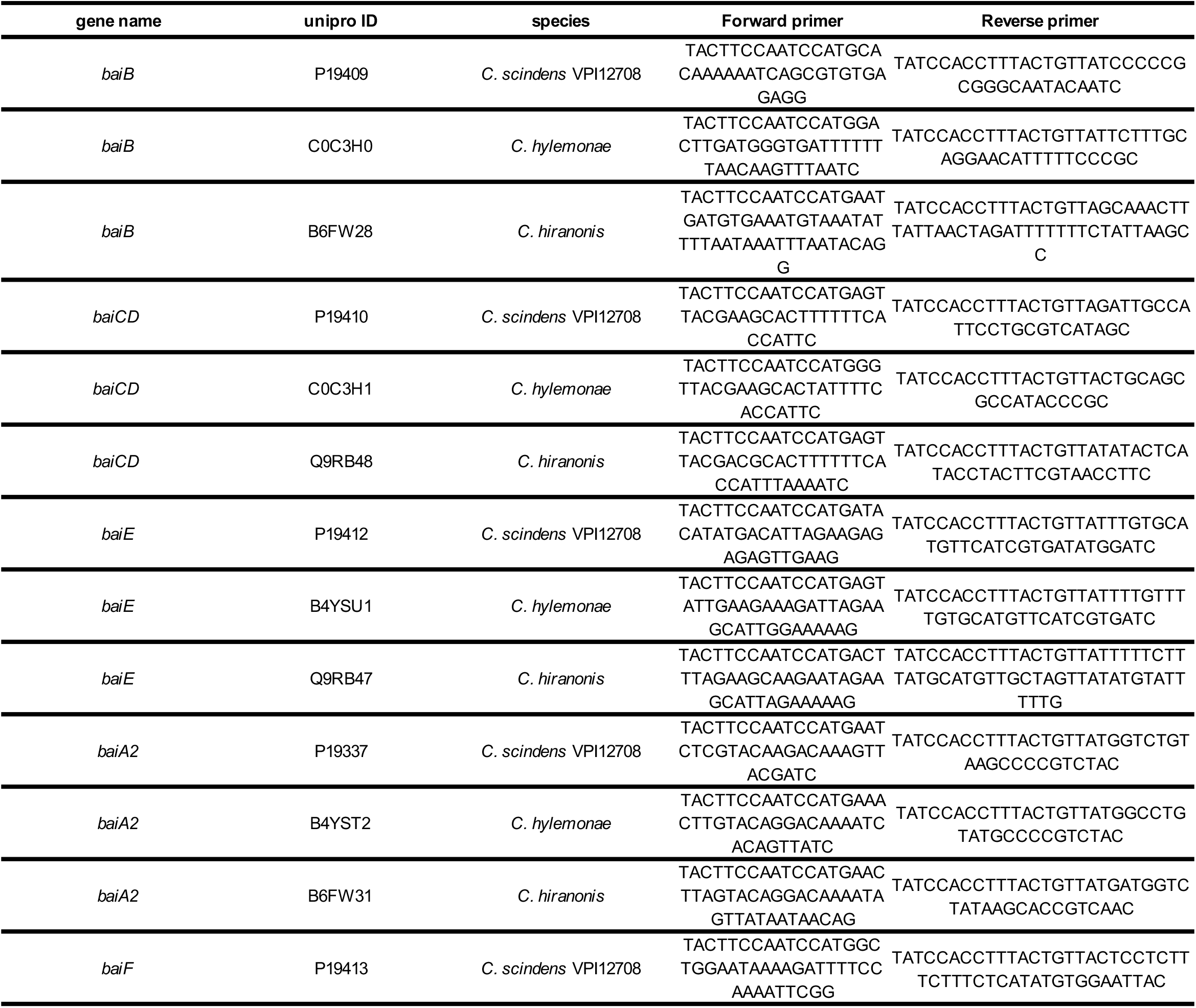
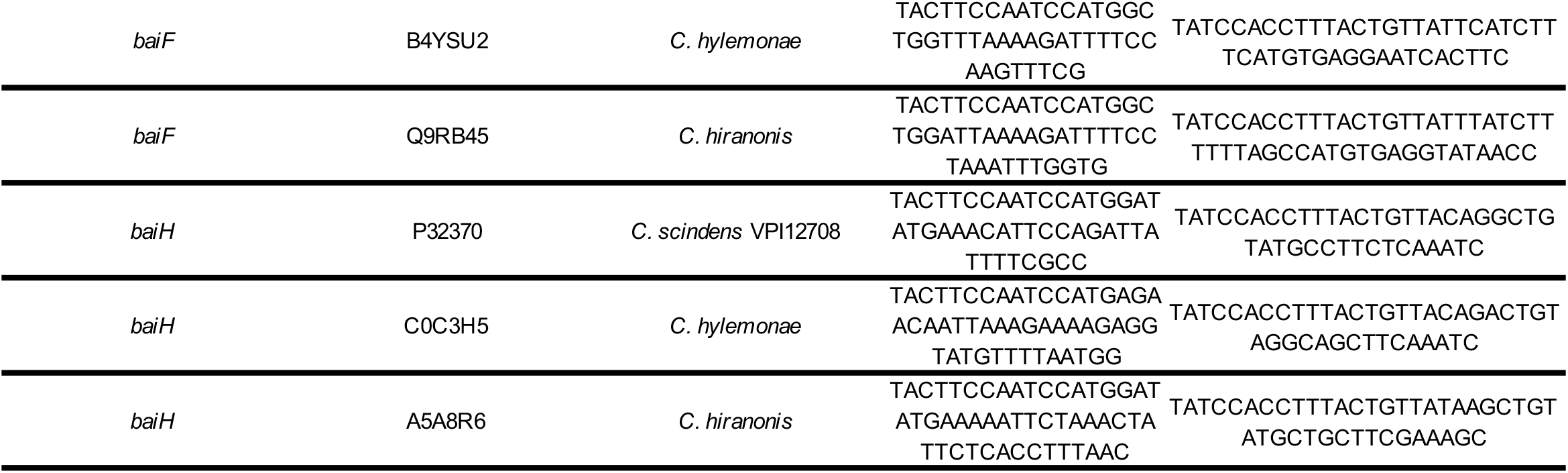
Primers used for expression of Bai proteins in *E. coli*

